# An isothermal calorimetry assay for determining steady state kinetic and enzyme inhibition parameters for SARS-CoV-2 3CL-protease

**DOI:** 10.1101/2024.01.31.578159

**Authors:** Luca Mazzei, Sofia Ranieri, Rebecca Greene-Cramer, Christopher Cioffi, Gaetano T. Montelione, Stefano Ciurli

**Affiliations:** Laboratory of Bioinorganic Chemistry, Department of Pharmacy and Biotechnology, University of Bologna, I-40127 Italy; Center for Biotechnology and Interdisciplinary Sciences, Rensselaer Polytechnic Institute, Troy, New York, 12180, USA; Department of Chemistry and Chemical Biology, Rensselaer Polytechnic Institute, Troy, New York, 12180, USA

## Abstract

This manuscript describes the application of Isothermal Titration Calorimetry (ITC) to characterize the kinetics of 3CL^pro^ from the Severe Acute Respiratory Syndrome CoronaVirus-2 (SARS-CoV-2) and its inhibition by Ensitrelvir, a known non-covalent inhibitor. 3CL^pro^ is the main protease that plays a crucial role of producing the whole array of proteins necessary for the viral infection that caused the spread of COVID-19, responsible for millions of deaths worldwide as well as global economic and healthcare crises in recent years. The proposed calorimetric method proved to have several advantages over the two types of enzymatic assays so far applied to this system, namely Förster Resonance Energy Transfer (FRET) and Liquid Chromatography-Mass Spectrometry (LC-MS). The developed ITC-based assay provided a rapid response to 3CL^pro^ activity, which was used to directly derive the kinetic enzymatic constants *K*_*M*_ and *k*_*cat*_ reliably and reproducibly, as well as their temperature dependence, from which the activation energy of the reaction was obtained for the first time. The assay further revealed the existence of two modes of inhibition of 3CL^pro^ by Ensitrelvir, namely a competitive mode as previously inferred by crystallography as well as an unprecedented uncompetitive mode, further yielding the respective inhibition constants with high precision. The calorimetric method described in this paper is thus proposed to be generally and widely used in the discovery and development of drugs targeting 3CL^pro^.

## INTRODUCTION

The Severe Acute Respiratory Syndrome CoronaVirus-2 (SARS-CoV-2) is the causal pathogen of coronavirus disease 2019 (COVID-19), which is responsible for more than seven million deaths worldwide and created a huge threat to the global economy and healthcare system ^1^. In the host infection by SARS-CoV-2, the two viral proteases 3CL^pro^ and PL^pro^ play the key role of catalyzing the hydrolysis of polyproteins pp1a and pp1ab, which are translated by host ribosomes upon recognition of the viral positive single-stranded RNA ^2,3^. The enzymatic cleavage of pp1a and pp1ab results in the release of an entire set of viral proteins crucial for SARS-CoV-2 replication ^4,5^.

SARS-CoV-2 3CL^pro^ is a homodimeric cysteine PA-clan protease ^6^, composed of two 33.8-kDa protomers, each consisting of three structural domains, namely domain I (residues 8 – 101), II (residues 102 – 184), and III (residues 201 – 306). The active site cleft, harboring the His41 - Cys145 (H41-C145) catalytic dyad (Figure 1), is located between domains I and II and consists of multiple subsites (S4, S3, S2, S1, and S1’) that, during catalysis, are filled by specific sequences of substrate amino acid residues (P4, P3, P2, P1, and P1’, respectively). The substrate recognition motif, which is highly conserved among several coronavirus 3CL^pro^, prefers the Leu-Gln-Ser (LQS) sequence in the P2-P1-P1’ position ^7^. 3CL^pro^ has little catalytic activity in its monomeric state and dimerization has been reported to be indispensable for a fully active protease ^8,9^. Crystallographic evidence suggests that the dimerization process involves the first seven residues at the N-terminus of each protomer (called the N-fingers), which appear to contribute to dimer stabilization and active site pocket architecture (the S1 subsite in particular) through interactions with domain II of the adjacent protomer as well as with domains II and III of the parent protomer ^8^. On the other hand, solution studies using native mass spectroscopy analysis support a model by which N-terminal processing is not critical for dimerization, which instead is triggered by induced fit due to covalent linkage during substrate processing ^10^.

**Figure 1.**
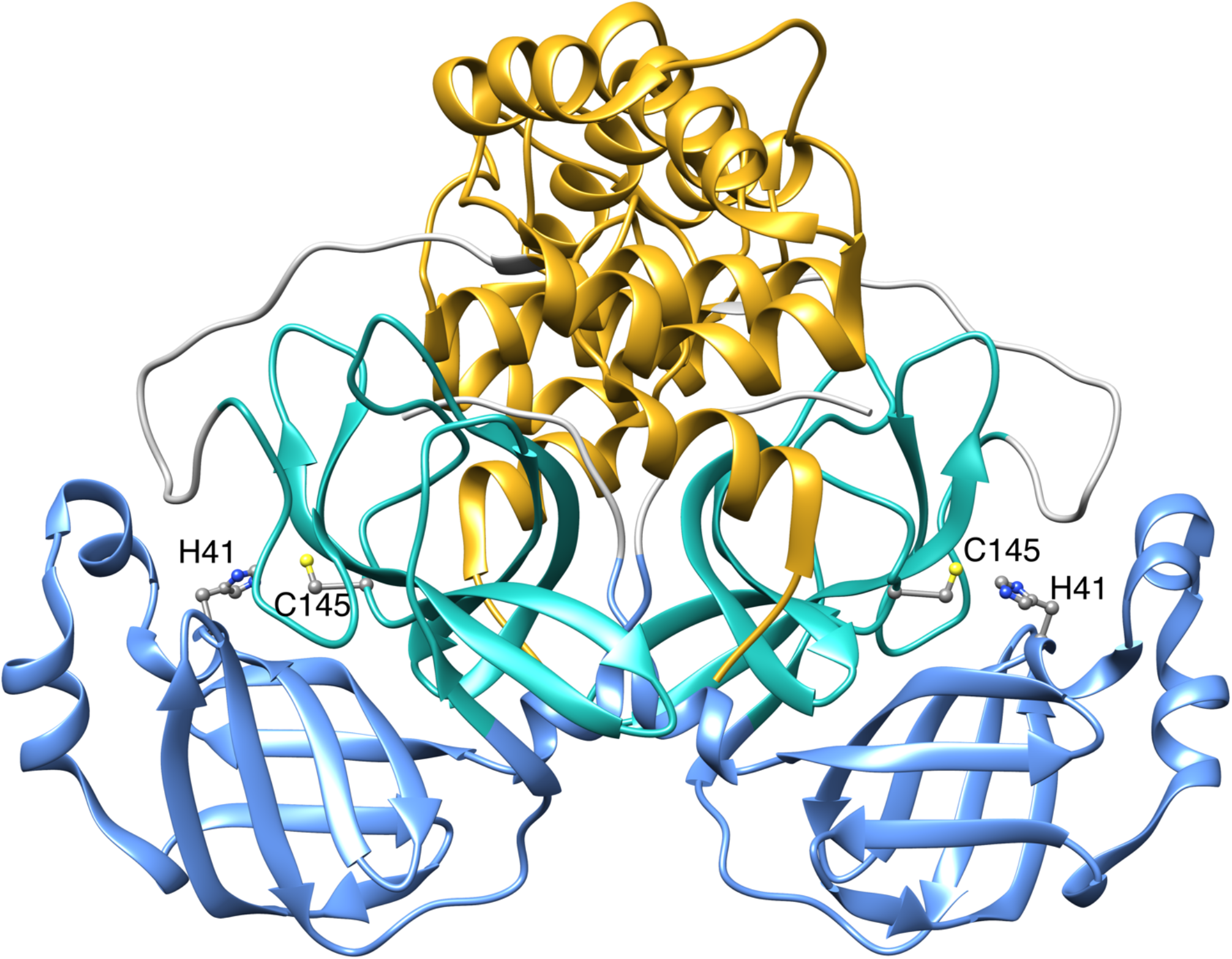
Ribbon representation of SARS-CoV-2 3CL^pro^ (PDB code 8CDC ^17^). Domains I, II, and III are colored in cornflower blue, light sea green and goldenrod, respectively. The position of the side chains of the catalytic dyad H41 and C145 are shown as ball-and-stick and colored according to the CPK code. Figure made using Chimera ^18^.

Given its essential role in the viral life cycle in conjunction with the absence of closely related homologs in the human genome, drug discovery targeting 3CL^pro^ has been a pivotal research area during the SARS-CoV-2 pandemic, and several 3CL^pro^ inhibitors have been identified ^11-14^. These molecules can be categorized into two large families, namely i) peptidomimetics, and ii) non-peptide small molecules. Peptidomimetics design is based on natural substrate scaffolds and typically share with the former the same binding subsites. They generally act as 3CL^pro^ inhibitors by exploiting an electrophilic warhead group (*i*.*e*., aldehydes, ketones, Michael acceptors, and nitriles) located near the P1 moiety to covalently bind the nucleophilic thiol group of C145. On the other hand, non-peptide small molecules reported to act as 3CL^pro^ inhibitors consist of members belonging to different families, including flavonoids, terpenoids, quinoline analogs, pyridinyl esters, Ebselen analogs, benzotriazole-based compounds, pyrimidine analogs, acrylamide and related compounds, isatin analogs, triazine compounds, and metal-containing analogs. Certain macrocyclic inhibitors of other PA-clan proteases are also inhibitors of the SARS CoV-2 3CL^pro 15,16^.

Despite the critical role played by 3CL^pro^ in the spread of COVID-19, detailed kinetic data on 3CL^pro^ activity and inhibition are limited, often presenting high variability depending on the methodology, on the form of 3CL^pro^ used in the assay, and on the different substrates used for the analysis ^19^. The two most common methods used to kinetically characterize 3CL^pro^ activity and inhibition are based on either <u>F</u>örster <u>R</u>esonance <u>E</u>nergy <u>T</u>ransfer (FRET) ^20-24^ or <u>L</u>iquid <u>C</u>hromatography-<u>M</u>ass <u>S</u>pectrometry (LC-MS) ^25,26^. The kinetic parameters *K*_*M*_ and *k*_*cat*_ obtained by the two techniques are often inconsistent and spanning several orders of magnitude: using FRET, values of *KM* in the 17 - 60 µM and 28 - 230 µM ranges were reported for 3CL^pro^ from SARS-CoV ^19,26,27^ and SARS-CoV-2 ^19,28,29^, respectively, while using LC-MS the corresponding values were 0.2 – 2.6 mM ^19,30^ and 0.9 mM ^19^. Concomitantly, the values reported for *kcat* by FRET are 0.2 – 2 s^-1^ and 0.05 – 0.23 s^-1^ for 3CL^pro^ from SARS-CoV ^19,26,27^ and SARS-CoV-2^19,28,29^, respectively, while LC-MS yields corresponding values of 0.54 – 6.4 s^-1 19,30^ or 2.2 s^-1 19^. Thus, the available values of *KM* estimated by LC-MS (in the mM range) are generally much higher than those reported for FRET-based methods (in the sub-millimolar range) while the values of *k*_*cat*_ span at least two orders of magnitude independently of the technique used in the assay. Furthermore, both FRET and LC-MS assays present important limitations: on one hand, FRET measurements can be affected by primary and secondary inner filter effects ^31^, substrate fluorophore interactions with the enzyme, and/or the absorbance or fluorescence of the inhibitor itself; conversely, while LC-MS has the advantage of being a label-free technique, it is also a complex and time-consuming sample manipulation procedure. Hence, there is a requirement for alternative approaches that can rapidly and effectively evaluate the catalytic and inhibitory efficiency of SARS-CoV-2 3CL^pro^ and its associated PA-clan proteases. This is essential for enhancing screening and optimization endeavors and gaining a deeper insight into the fundamental mechanisms of enzyme inhibition.

Isothermal <u>T</u>itration <u>C</u>alorimetry (ITC) is a technique that can be applied to characterize enzyme kinetics by monitoring the time-course of the heat generated upon rapid mixing of a series of small-volume injections of a substrate (or an enzyme) solution from a computer-controlled syringe into a sample cell containing an enzyme (or substrate) solution, in the absence or presence of different concentrations of a given inhibitor ^32-38^. This approach is very versatile, as most chemical reactions involve the production or consumption of heat. Unlike other techniques inferring catalysis rates indirectly from substrate and/or product concentrations, ITC provides a real-time detection of heat flow, offering a direct readout of enzyme activity and its modulation in response to inhibitors. ITC does not require the development of customized assays using fluorophores or chromophores as substrates or products as for FRET, nor post-reaction separation of products and substrates as for LC-MS. Moreover, in comparison to standard spectroscopic measurements, where enzyme, substrate, and inhibitor solutions are combined with delays before the start of the measurement, ITC measures heat flow rapidly, minimizing dead time. To the best of our knowledge, despite its considerable potential, there has been no study to date that has utilized ITC to characterize the kinetics of catalysis and inhibition of 3CL^pro^.

Here we present a full characterization of the activity of 3CL^pro^ from SARS-CoV-2 carried out using ITC methods that provide kinetic parameters in less than an hour, with high reproducibility. The developed assay is also applied to study the inhibition of 3CL^pro^ by the well-established 3CL^pro^ inhibitor Ensitrelvir, the first oral non-covalent and non-peptide inhibitor (belonging to the triazine compounds) developed by Shionogi ^39^ which was approved for emergency use in Japan in November 2022 (sold under the brand name Xocova, Scheme 1).

**Scheme 1:**
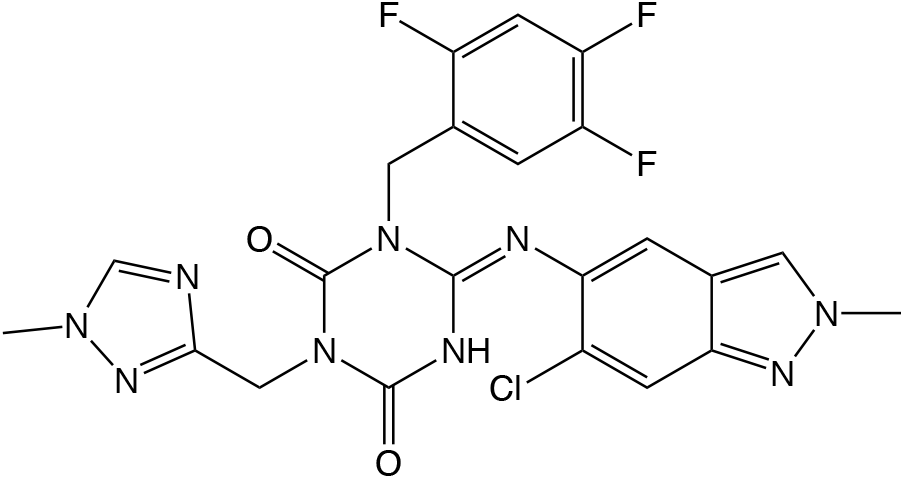
chemical structure of Ensitrelvir

## RESULTS

### Enzyme kinetics

Classically, any enzyme catalyzed reaction could best be described by the following reaction Scheme 2, where E is the enzyme, S is the substrate and P is the product of the reaction:

**Scheme 2:**
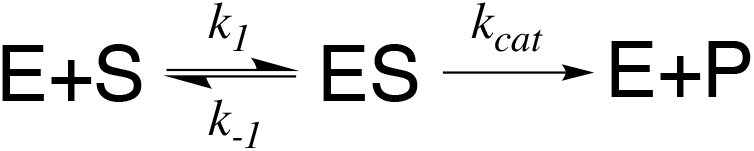
classical enzyme kinetics

In the case of 3CL^pro^, the first step is the reversible binding of the enzyme to its substrate with the formation of a stable ES complex, governed by the Michaelis constant *K*_*M*_, which is the equilibrium dissociation constant (*k*_*-1*_*/k*_*1*_) of the ES complex and represents the concentration of substrate required to achieve a half-maximal reaction rate; the second step involves the activation of the ES complex and its subsequent decomposition into products and free enzyme and is governed by *k*_*cat*_ (s^-1^), the catalytic rate constant (also known as turnover number) that describes the limiting number of chemical conversions of substrate molecules per second.

The most used approach to mathematically treat Scheme 2 is the Michaelis–Menten model, which describes the reaction rate, expressed as the time-dependent decrease of substrate concentration [S], as a function of [S] and [E], as well as a function of *K*_*M*_ and *k*_*cat*_:

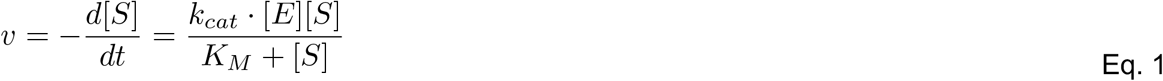

Here, *k*_*cat*_ •[E], also defined as *V*_*max*_, is the maximum reaction rate in the theoretical presence of an infinite amount of substrate. From the Michaelis-Menten equation, it is clear that the determination of *K*_*M*_ and *k*_*cat*_ for an enzyme-catalyzed reaction described by this simple model provides its complete characterization.

### Principles of ITC methodology

An isothermal titration calorimeter (Figure 2A) consists of an adiabatic shield encompassing two cells, namely a *reference cell* (generally filled with deionized water) and a *sample cell*, both having openings to the outside for solutions introduction or removal *via* long-needled syringes. A rotating paddle-shaped syringe is mounted on the sample cell, where it dispenses its content and provides complete mixing of the two solutions after an injection.

**Figure 2.**
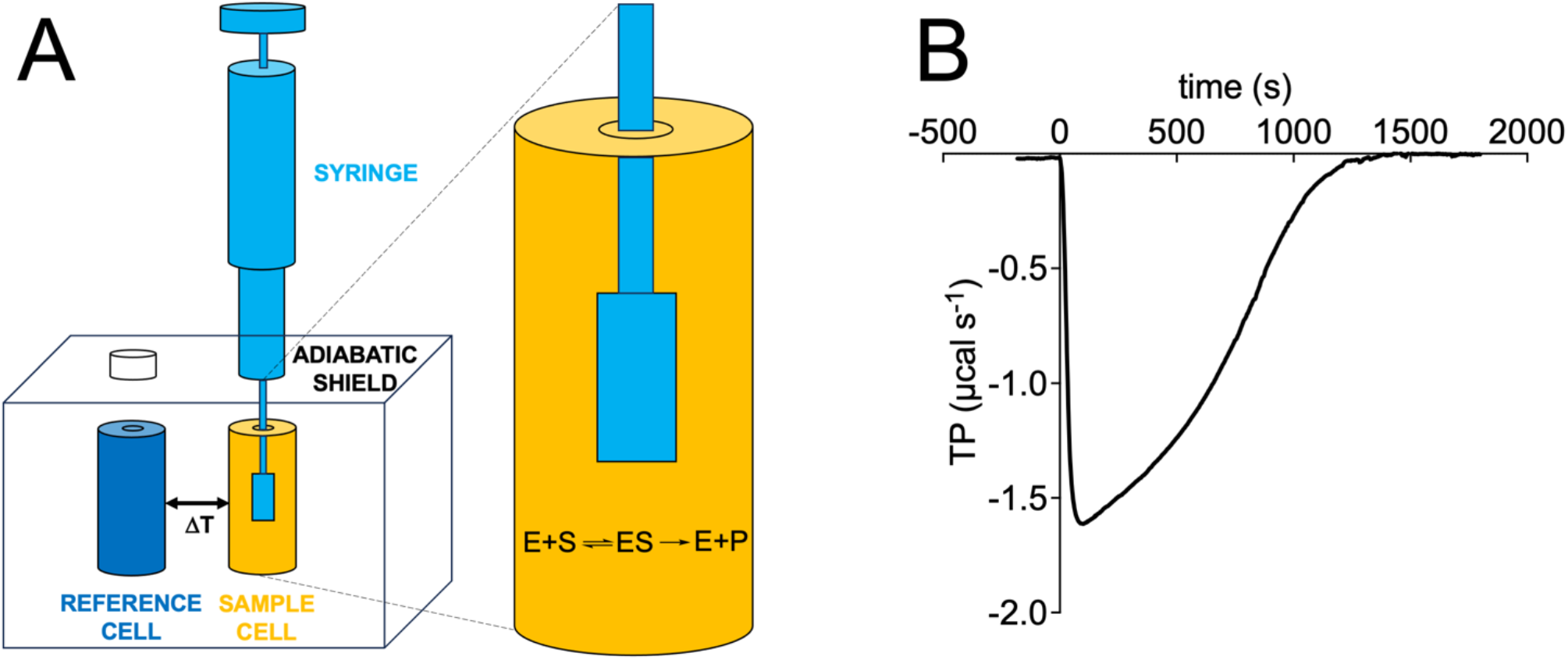
ITC instrumentation. (**A**) Schematic representation of the isothermal titration calorimeter: reference and sample cells are indicated in blue and orange, respectively, and the titration syringe is colored in light blue. (**B**) ITC output, provided as thermal power required to maintain constant temperature after addition of substrate to enzyme (or *vice versa*), is represented as a deviation of the baseline in the recorded data.

During an ITC experiment, a thermoelectric device measures in real time the difference in temperature between the sample and the reference cell and, using a cell feedback network, it maintains this difference (ΔT) at zero by adding/removing heat to/from the sample cell, the latter being recorded over the time course of the experiment. The amount of heat (*Q*) provided or removed by the system as a function of time (*t*) is defined as the thermal power (TP, often also referred to as heat flow) (Eq. 2, Figure 2B):

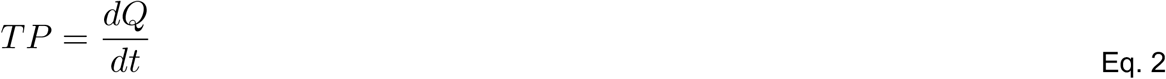

In any enzyme-catalyzed reaction, the amount of heat associated with the conversion of *n* moles of substrate to product is given by Eq. 3:

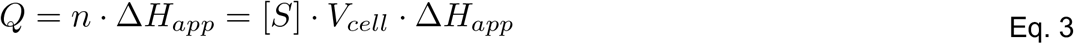

where *ΔH*_*app*_ is the total apparent molar enthalpy for the reaction under study, [S] is the molar concentration of converted substrate, and *V*_*cell*_ is the volume of the sample cell where the reaction occurs. The reaction rate, defined as the change in substrate concentration over time, can thus be related to the thermal power as given by Eq. 4:

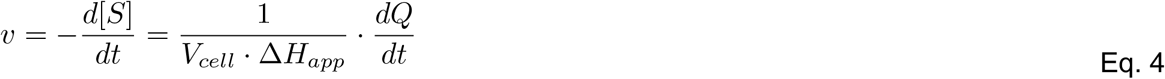

From Eq. 4 it becomes clear that in order to calculate the reaction rates as a function of the substrate concentration, which then can be fitted using the Michaelis-Menten equation (Eq. 1) to derive the kinetic parameters *K*_*M*_ and *k*_*cat*_, it is necessary to (i) know the total apparent molar enthalpy *ΔH*_*app*_ and (ii) measure *dQ/dt* at different substrate concentrations.

*ΔH*_*app*_ is usually derived by means of the so-called *direct single-injection* method: a small concentration of substrate (below the expected value of *K*_*M*_) is injected into a solution containing the enzyme (usually in the nM–μM range). The thermal power generated by the reaction is measured over time until the substrate is completely consumed and the signal returns to the pre-injection baseline; the concentrations of enzyme and substrate are chosen so that complete conversion occurs on the timescale of minutes or tens of minutes. Integration of the area under the curve yields the experimental *ΔH*_*app*_ according to Eq. 5:

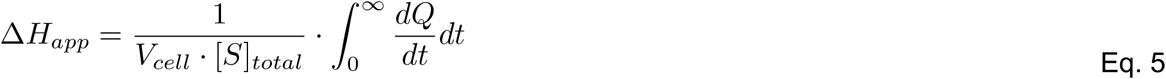

where [S]_total_ is the total concentration of substrate present in the sample cell at the beginning of the experiment.

The *dQ/dt* values at different substrate concentrations are usually determined by carrying out the so-called *multiple-injection* method: subsequent small injections of a concentrated substrate solution in the syringe are performed into a diluted enzyme solution (in the pM–nM range). Each injection provides an increase of substrate concentration, which is reflected in a displacement of the baseline that is in turn indicative of a change of thermal power in the sample cell. The injections are spaced in time so that the thermal power can stabilize at the new baseline level but must be short enough to avoid the conversion of a significant amount of substrate (less than 5 %) thus allowing the measurement to be performed under steady-state conditions. The value of *dQ/dt* at each substrate concentration is determined by measuring the difference between the original baseline and the new baseline after each injection. The resulting reaction rates as a function of the substrate concentration can thus be calculated using Eq. 4 and fitted to the Michaelis-Menten curve (Eq. 1) to derive the kinetic parameters *K*_*M*_ and *k*_*cat*_. The main disadvantage of this *multiple-injection* method is that the total generated thermal power is often quite low, especially if the reaction under study features a small *ΔH*_*app*_; this situation could render the assay sensitive to baseline drifts and instrumental noise, which in turn could affect the reliability of *K*_*M*_ and *k*_*cat*_ estimates. Concomitantly, the required high concentrations of substrate in the syringe solution might be difficult to attain for sparingly soluble compounds, while the large heat of substrate dilution would significantly interfere with the measurement of the heat of reaction.

This is the situation that was encountered while investigating the enzymatic activity of 3CL^pro^, thus prompting us to alternatively investigate the so-called *inverse single-injection* experiment, in which a small amount of enzyme solution (in the order of tens of µM) is injected from the syringe into the sample cell containing a substrate solution at a concentration much higher than the expected *K*_*M*_ (but not necessarily as high as to incur into solubility issues) so that the injection significantly saturates the enzyme. This setup can also be used to study the enzyme inhibition kinetics by adding known concentrations of inhibitors to the substrate solution in the sample cell.

In this experiment, a large *dQ/dt* deflection occurs immediately after the injection of the enzyme solution because of the heat released by the reaction (Figure 3A), reaching a maximal effect that gradually decreases as the substrate concentration in the sample cell is reduced due to the enzyme-catalyzed reaction, eventually returning to the pre-injection baseline as the substrate is fully consumed. The variation of substrate concentration in any time interval (that corresponds to 2 seconds according to the ITC data output) can be determined from Eq. 6:

**Figure 3.**
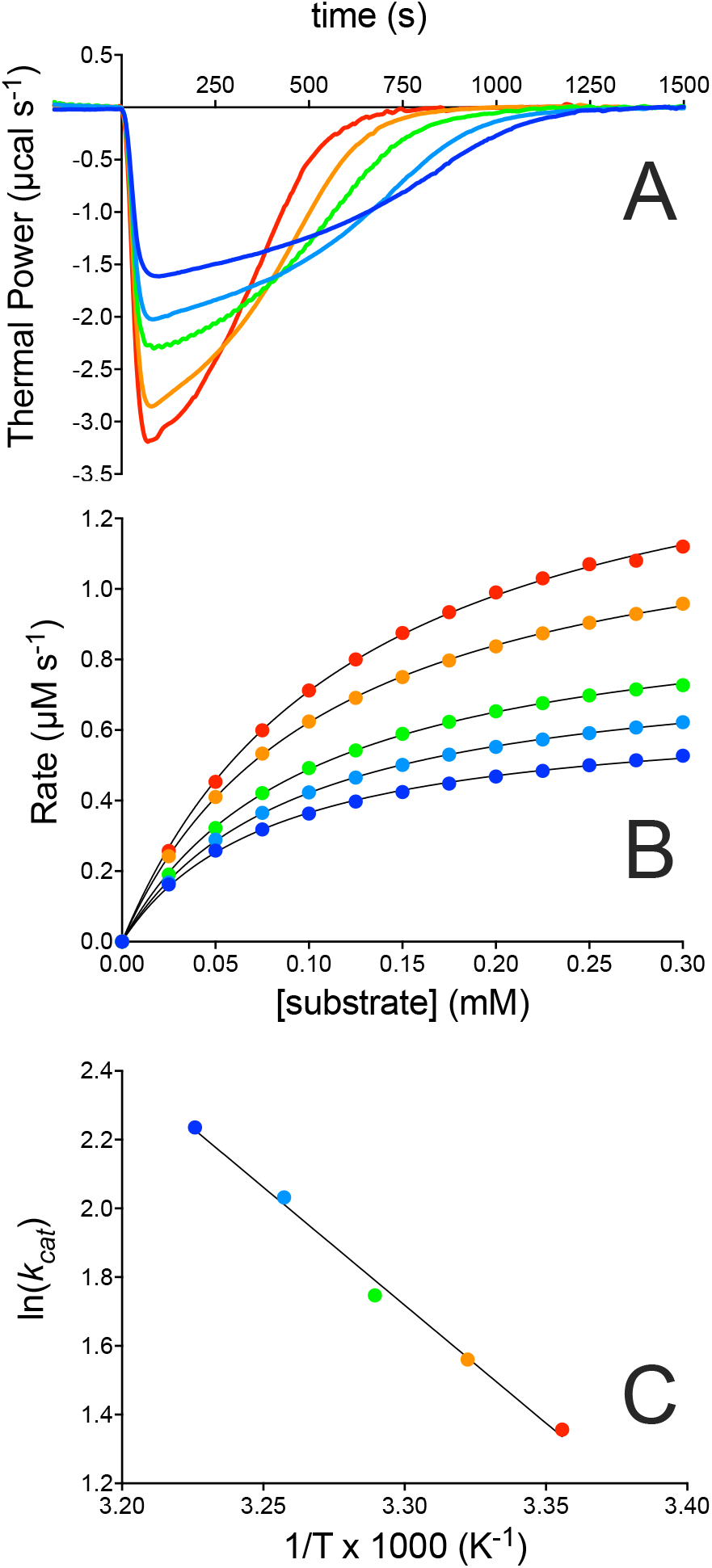
3CL^pro^ kinetics at pH 7.5 characterized by ITC. (**A**) Thermal power recorded as a function of time after the injection of 150 nM 3CL^pro^ in the sample cell containing 350 µM substrate at 298 K (black line), 301 K (blue line), 304 K (green line), 307 K (orange line), and 310 K (red line). (**B**) Reaction rates calculated from the raw data shown in **A** using Eq. 4. Data fits using the Michaelis-Menten equation are also shown as colored lines. (**C**) Arrhenius plot obtained by plotting the logarithm of the *k*_*cat*_ values derived at five different temperature values versus the inverse temperature, 1/T.

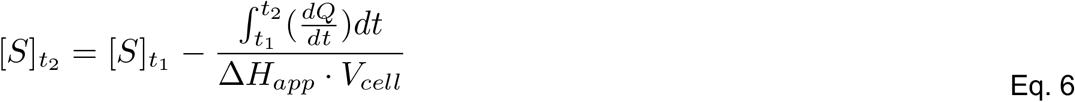

Here, *ΔH*_*app*_ is directly obtained by integration of the full area under the *TP vs*. time curve, according to Eq. 5, while *t*_*1*_ and *t*_*2*_ are two consecutive time points. The reaction rates as a function of substrate concentration can be derived using Eq. 4 and then fitted using the Michaelis-Menten equation (Eq. 1) to derive *K*_*M*_ and *k*_*cat*_. The reaction rates shown in Figure 3B represent a subset of the experimentally measured data points (in the order of a few hundreds), arbitrarily selected every 0.025 mM for the sake of clarity.

### Derivation of the kinetic parameters of 3CL^pro^ enzymatic catalysis

The kinetics of 3CL^pro^ was studied at five different temperatures between 298 and 310 K (Figure 3). The calorimetric raw data are presented in Figure 3A and show, in all cases, a large decrease of the thermal power immediately after the injection, depicting an exothermic reaction event that eventually returns to the pre-injection baseline as the substrate is completely consumed. The heat of mixing, measured in a separate experiment by injecting the enzyme solution onto buffer alone, was negligible. Integration of the raw data yielded a value of *ΔH*_*app*_ *ca*. -2 kcal mol^-1^, consistent with values previously reported for the hydrolysis of a peptide bond in the same buffer and temperature conditions ^40^.

The reaction rates and corresponding fits carried out using the Michaelis-Menten equation are shown in Figure 3B. The values of *K*_*M*_ = 81 ± 2 µM and *k*_*cat*_ = 3.9 ± 0.1 s^-1^ were obtained at 298 K, while the full series of thermodynamic and kinetic parameters derived from the experiments conducted at increasing temperatures are reported in Table 1. These data reveal an effective invariance of *ΔH*_*app*_ with respect to temperature, a slight increase of *K*_*M*_ from 81 to 124 µM, which indicates a small reduction in the enzyme affinity for the substrate at higher temperature, as well as an expected increase of *k*_*cat*_ in the 298 – 310 K range. The dependence of *k*_*cat*_ as a function of the temperature provided the value of the activation energy (*E*_*a*_ = 13.1 ± 1.4 kcal mol^-1^) of the 3CL^pro^ hydrolytic reaction, derived from the Arrhenius equation (Eq. 7) and the corresponding Eyring plot (Figure 3C):

**Table 1.** Kinetic parameters of 3CL^pro^ measured at different temperatures

| | $\Delta H_{app}$<br>(kcal mol<sup>-1</sup>) | $K_M$ (μM) | $k_{cat}$<br>(s<sup>-1</sup>) | $k_{cat}/K_M$ (M<sup>-1</sup> s<sup>-1</sup>) |
| --- | --- | --- | --- | --- |
| **298 K** | 2.08 | 81 ± 2 | 3.9 ± 0.1 | 48 ± 2.4 |
| **301 K** | 2.16 | 92 ± 1 | 4.8 ± 0.1 | 52 ± 1.7 |
| **304 K** | 2.20 | 100 ± 1 | 5.7 ± 0.1 | 57 ± 1.6 |
| **307 K** | 2.01 | 109 ± 1 | 7.6 ± 0.1 | 70 ± 1.6 |
| **310 K** | 1.97 | 124 ± 3 | 9.3 ± 0.1 | 75 ± 2.6 |

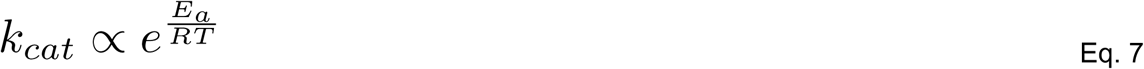

Substrate or product inhibition in this experimental setup were ruled out through the analysis of the shape of the obtained Michaelis-Menten curves as well as by carrying out experiments with different substrate concentrations in the 150 – 350 µM range and observing that the values of *K*_*M*_ and *k*_*cat*_ are not significantly affected.

### Derivation of the kinetic parameters for the inhibition of 3CL^pro^ with Ensitrelvir

In addition to the kinetic characterization of the enzymatic 3CL^pro^ reaction, *inverse single-injection* experiments were also used to carry out a kinetic characterization of 3CL^pro^ inhibition by Ensitrelvir (Figure 4), whereby an enzyme solution was injected into the sample cell containing the substrate and increasing concentrations of inhibitor.

**Figure 4.**
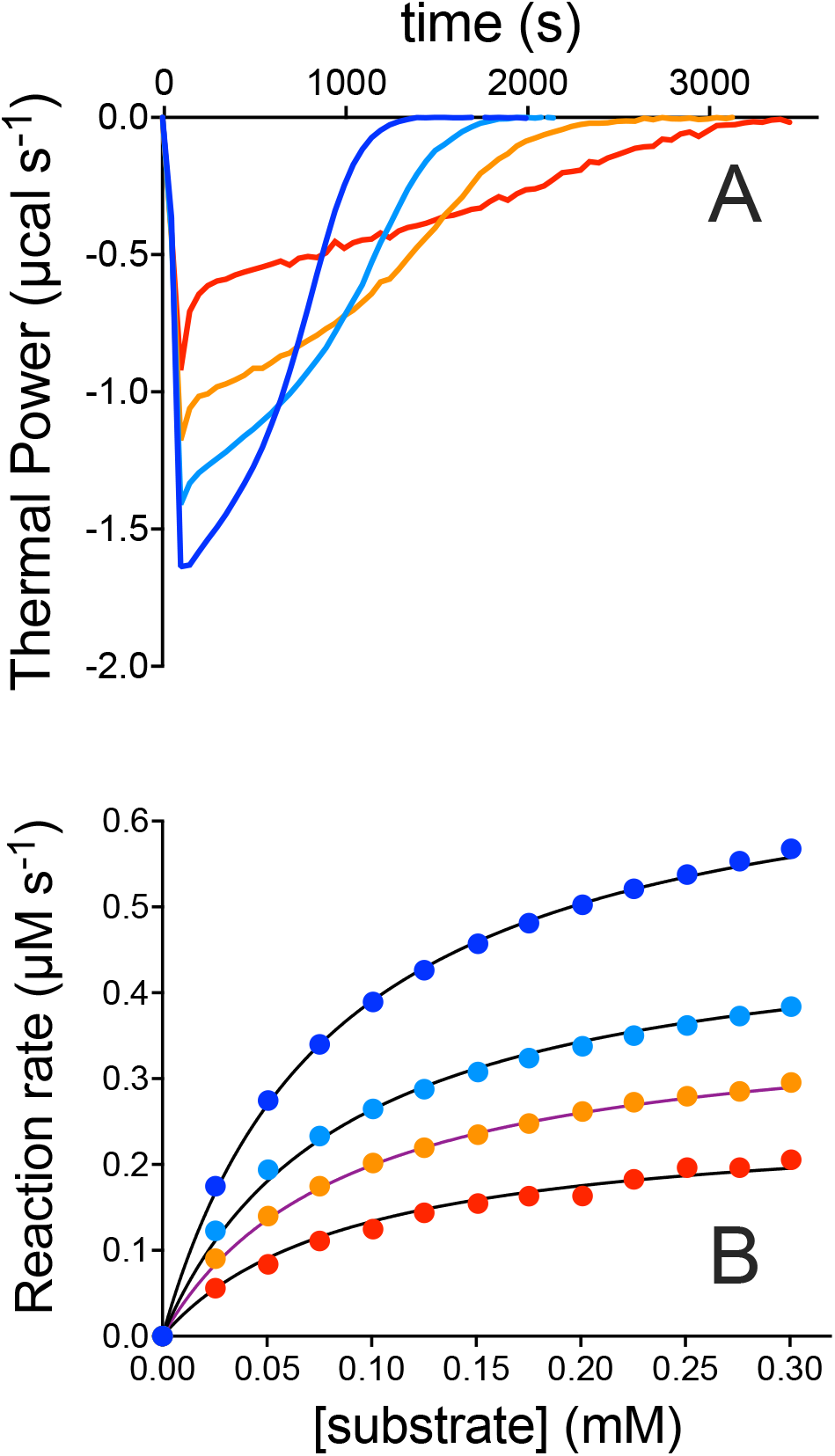
3CL^pro^ inhibition by Ensitrelvir at pH 7.5 characterized by ITC. (**A**) Thermal power recorded as a function of time after the injection of 150 nM 3CL^pro^ in the sample cell containing 350 µM substrate, also containing 25 nM (blue line), 50 nM (pink line), and 100 nM (red line). (**B**) Reaction rates calculated from the raw data shown in **A** using Eq. 4 and shown colored according to **A**. Data fitting using the Michaelis-Menten equation are shown as solid lines.

Raw data recorded from the experiments performed at 298 K in the presence of 25 nM, 50 nM, and 100 nM Ensitrelvir are shown in Figure 4A. The time needed for the signal to return to the pre-injection baseline level progressively increased at increasing inhibitor concentration; however, integration of the raw data provided an averaged *ΔH*_*app*_ of *ca*. -2 kcal mol^-1^, revealing that complete substrate consumption occurs over longer time periods by increasing [I] (*i*.*e*., the presence of the inhibitor at the tested concentrations slows down the hydrolytic activity of the enzyme but does not prevent it completely). The reaction rates of 3CL^pro^ in the presence of the three Ensitrelvir concentrations were globally fit, together with the data calculated from the non-inhibited 3CL^pro^, using Eq. 7:

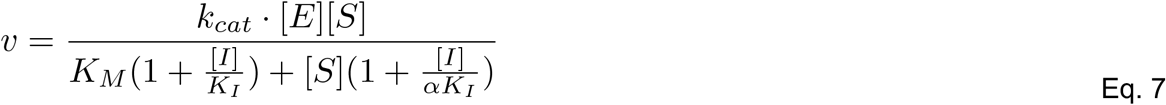

In this equation, [I] corresponds to the chosen inhibitor concentrations, *K*_*I*_ is the competitive inhibition constant, and the α value describes whether the type of inhibition is competitive (α >> 1), uncompetitive (α << 1), or non-competitive (α = 1), with α*K*_*I*_ corresponding to the uncompetitive inhibition constant ^41^. The fit provided a value of *K*_*I*_ = 46 ± 3 nM with α = 1.2 ± 0.1, consistent with a non-competitive inhibition mechanism for Ensitrelvir ^41^.

## DISCUSSION

This study represents the first successful application of isothermal titration calorimetry to the characterization of the enzymatic activity of SARS-CoV-2 3CL^pro^ and its inhibition by Ensitrelvir, a non-covalent inhibitor. The technique has proven to be highly effective, rapid, reproducible, reliable, and relatively easy to apply. Using ITC, Michaelis-Menten profiles can be obtained in under 1 hr. The exploration of different modes of applications such as *direct multiple-injections* of substrate into an enzyme solution, as well as *inverse single-injection* of enzyme into a substrate solution in the absence or presence of different concentrations of the inhibitor allowed us to identify in the latter experimental setup the optimal conditions to quickly extract the kinetic and inhibition parameters with high precision. The approach using ITC in this *inverse single-injection* mode has several advantages with respect to the types of assays previously employed for the characterization of kinetics and inhibition of 3CL^pro^, namely FRET or LC-MS. The most important aspect is that calorimetry continuously measures enzyme catalytic rates without relying on complicated procedures to measure substrate or product concentrations in batch experiments. As a result, data are effectively collected for hundreds of different substrate concentrations which allows for the determination of a highly robust Michaelis-Menten profiles. In addition, no specific probes are needed, nor post-reaction sample manipulations are employed. The versatility of this technique is proven by its capability to provide the catalytic parameters of the enzymatic reaction, such as *K*_*M*_ and *k*_*cat*_ in less than 1 hr.

The obtained kinetic parameters for the specific peptide hydrolysis catalyzed by SARS-CoV-2 3CL^pro^ are consistent with previous values obtained by FRET in the case of *K*_*M*_ (81 ± 2 µM) ^19,28^, while *k*_*cat*_ (3.9 ± 0.1 s^-1^) is more like values obtained by LC-MS ^19^.

The ITC approach also revealed that Ensitrelvir inhibits SARS-CoV-2 3CL^pro^ in a non-competitive mode, directly providing the values of the competitive (46 ± 3 nM) and uncompetitive (55 ± 4 nM) inhibition constants. So far, neither FRET nor LC-MS have been used to establish the mode of inhibition of Ensitrelvir, which has been *considered* competitive only based on the X-ray structures of the enzyme-inhibitor complex in the solid state ^39,42,43^. Moreover. neither FRET nor LC-MS, only giving IC_50_ values, have thus far directly provided values for the inhibition constant; indeed the only reported value of *K*_*I*_ = 9 ± 0.7 nM was calculated from the IC_50_ value *assuming* a competitive mechanism ^43^. The range of IC_50_ values (13 - 49 nM) reported for Ensitrelvir and determined using FRET ^43-45^ or LC-MS ^39^ are in the same order of magnitude of the inhibition constants measured by ITC.

The non-competitive mode of inhibition of SARS-CoV-2 3CL^pro^ protease by Ensitrelvir, established by ITC, is an unprecedented result. Ensitrelvir appears to effectively operate by binding not only to the empty active site cavity preventing the concomitant binding and processing of the substrate, as revealed by crystallography, but also to the pre-formed ES complex with similar affinity. It is important to highlight that while the assays based on FRET or LC-MS have entailed a wide range of enzyme-inhibitor incubation times (1.5 – 180 min), during which a stable complex can form prior to its exposure to the substrate, the described calorimetric assay is carried out by injecting the enzyme in a solution that contains both substrate and Inhibitor, thus allowing a true competition.

The developed calorimetric assay allowed us to also determine the temperature-dependence of the catalysis, with the obtainment of the activation energy for the reaction. The value of *E*_*a*_ = 13.1 ± 1.4 kcal mol^-1^ is, to the best of our knowledge, the first experimentally obtained activation energy for the 3CL^pro^ catalytic hydrolysis. This value agrees with previous theoretical studies on 3CL^pro 46,47^ as well as with reported experimental and computational data for other cysteine proteases ^48^.

## CONCLUSIONS

We propose the generalized use of the calorimetric assay developed in this study to investigate the kinetics of catalysis and inhibition of SARS-CoV-2 3CL^pro^. ITC can also be used to validate novel hits from a 3CL^pro^ inhibitor high-throughput (HT) screen and eliminate potential pan-assay interference compounds (PAINS) that can provide false positives in widely used HT fluorescence-based platforms ^49^. Moreover, in addition to the determination of inhibition modes and relative constants, ITC can provide critical thermodynamic signatures such as binding enthalpy and entropy for protein-ligand binding that can assist with drug design efforts toward improved 3CL^pro^ inhibitors. Indeed, focus on enhanced enthalpic *vs*. entropic contributions to binding for hit compounds is considered favorable and has been reported to lead to better prioritization and optimization of ligands for potential hit selection and hit-to-lead optimization ^49,50^.

## MATERIALS AND METHODS

### Enzyme and inhibitor sources

Native 3CL^pro^ (M_r_ monomer = 33.8 kDa, pI = 5.95) was expressed and purified following a previously described protocol ^17^. The protein (purity > 98% as checked by SDS-PAGE) was stored as 150 µM stock aliquots (protein concentration is referred to the monomer throughout the manuscript) at -80 °C in 20 mM Tris-HCl buffer, 50 mM NaCl, at pH 7.5. The peptide substrate WKTSAVLQ/SGFRKMEW (M_r_ = 1.95 kDa, pI = 9.99) was designed with the 3CL^pro^ cleavage site Q/S, and was purchased from GenScript (Rijswijk, Netherlands). It includes two non-native tryptophan (W) residues at the N and C termini to ensure a measurable absorption at 280 nm. Solutions of substrate were freshly prepared before every experiment (see below for details). Protein and peptide quantification was carried out by measuring the absorbance at 280 nm considering a molar extinction coefficient (ε_280_) of 33,000 M^-1^ cm^-1^ and 11,000 M^-1^ cm^-1^ for 3CL^pro^ and the substrate respectively, estimated using ProtParam ^51^. Ensitrelvir fumarate was purchased from Cabru S.A.S. (Arcore, Italy), dissolved at 10 mM in pure DMSO, and stored as 10 µL aliquots at -80 °C.

### Calorimetric studies on the enzymatic hydrolysis by 3CL^pro^

The determination of the 3CL^pro^ kinetic parameters was carried out using a high-sensitivity VP-ITC micro-calorimeter (MicroCal LLC, Northampton, MA, USA). For each experiment, the reference cell was filled with deionized water and the temperature of the reference and sample cells was set and stabilized at five temperatures in the range 298 - 310 K. Stirring speed was 300 rpm and the thermal power was monitored every 2 s using high instrumental feedback. Solutions of 3CL^pro^ for the assay were prepared by diluting the stock solution down to 1.5 µM enzyme concentration in 600 µL of 20 mM Tris-HCl, 50 mM NaCl, 1 % DMSO, at pH 7.5 and loaded into the injection syringe. The measured enzymatic activity was not affected over several days at 4 °C in the absence of DTT. The substrate was prepared by dissolving 2 mg of the purchased powder in 2 mL of the same buffer, obtaining a final concentration of 350 µM, and loaded in the sample cell (final volume = 1.50 mL). Accurate protein and substrate quantification was carried out, as described above, prior to each experiment. The *inverse single-injection* experiment was carried out by injecting 15 µL of the 1.5 µM 3CL^pro^ solution (final enzyme concentration in the sample cell = 0.15 µM) from the syringe into the 350 µM substrate solution. The thermal power (TP, µcal s^-1^) was recorded over a time of 2000 s, thus ensuring the instrument baseline shift caused by the heat flow generated from the enzymatic hydrolysis of the substrate, to return to its original pre-injection level. The raw calorimetric data were processed using the MicroCal Origin software to derive the total apparent molar enthalpy *ΔH*_*app*_ of 3CL^pro^-catalyzed substrate hydrolysis (according to Eq. 5), as well as the reaction rates as a function of substrate concentration (according to Eqs. 4 and 6). The obtained reaction rates were fit using the canonical Michaelis-Menten equation (Eq. 1) to derive the kinetic parameters *K*_*M*_ and *k*_*cat*_.

### Calorimetric studies on the enzymatic inhibition of 3CL^pro^ by Ensitrelvir

The same *inverse single-injection* setup was used to perform three additional experiments at 298 K where the 15 µL injection of the 1.5 µM 3CL^pro^ solution was carried out into the 350 µM substrate solution also containing increasing concentrations of Ensitrelvir (in the range 25 – 100 nM). The thermal power was continuously recorded over a variable experimental time (in the range 2000 – 4000 s, depending on the inhibitor concentration) to ensure the return of the instrument baseline to its original pre-injection level. Calorimetric raw data were processed as described above, and the resulting reaction rates as a function of substrate concentration, calculated for each concentration of inhibitor, were simultaneously and globally fit, together with the data calculated from the non-inhibited 3CL^pro^, using Eq. 7 to determine the inhibition constant (*K*_*I*_) and the mode of action (through the α value) of Ensitrelvir for 3CL^pro^.

## ACKNOWLEDGMENTS

LM and SC acknowledge financial support from the University of Bologna, the Consorzio Interuniversitario di Risonanze Magnetiche di Metallo-Proteine (CIRMMP). RG-C was supported by a NIGMS Graduate Training Grant (T32-GM141865). GTM was supported by grant R35-GM141818 (to GTM) from the National Institutes of Health, National Institute of General Medical Sciences.

